# Essential Tremor Disrupts Rhythmic Brain Networks During Naturalistic Movement

**DOI:** 10.1101/2024.06.26.600740

**Authors:** Timothy O. West, Kenan Steidel, Tjalda Flessner, Alexander Calvano, Deniz Kucukahmetler, Marielle J. Stam, Meaghan E. Spedden, Benedikt Wahl, Veikko Jousmäki, John Eraifej, Ashwini Oswal, Tabish A. Saifee, Gareth Barnes, Simon F. Farmer, David J. Pedrosa, Hayriye Cagnan

## Abstract

Essential Tremor (ET) is a very common neurological disorder characterised by involuntary rhythmic movements attributable to pathological synchronization within corticothalamic circuits. Previous work has focused on tremor in isolation, overlooking broader disturbances to motor control during naturalistic movements such as reaching. We hypothesised that ET disrupts the sequential engagement of large-scale rhythmic brain networks, leading to both tremor and deficits in motor planning and execution. To test this, we performed whole-head neuroimaging during an upper-limb reaching task using high-density electroencephalography in ET patients and healthy controls, alongside optically pumped magnetoencephalography in a smaller cohort. Key motor regions—including the supplementary motor area, premotor cortex, posterior parietal cortex, and motor cerebellum—were synchronized to tremor rhythms. Patients exhibited a 15% increase in low beta (14-21 Hz) desynchronization over the supplementary motor area during movement, which strongly correlating with tremor severity (R^2^ = 0.85). A novel dimensionality reduction technique revealed four distinct networks accounting for 97% of the variance in motor-related brain-wide oscillations, with ET altering their sequential engagement. Consistent with our hypothesis, the frontoparietal beta network- normally involved in motor planning-exhibited additional desynchronization during movement execution in ET patients. This altered engagement correlated with slower movement velocities, suggesting an adaptation towards feedback-driven motor control. These findings reveal fundamental disruptions in distributed motor control networks in ET and identify novel biomarkers as targets for next-generation brain stimulation therapies.

## 1 Introduction

The ability to effectively control movement is a primary function of the nervous system [1] and is critically impaired in neurological disorders such as Essential Tremor (ET). Many neurological disorders are associated with hypersynchronous oscillations [2], that can disrupt activity in neural circuits and lead to debilitating symptoms. In ET, pathological synchrony manifests as involuntary rhythmic tremors affecting the limbs, head, or trunk [3]. A wealth of neuroimaging work supports the existence of a distributed central tremor circuit [4,5], involving the thalamus, cerebellum, parietal, and motor cortex. This network has been established using both hemodynamic correlates of tremor [6–9], as well as electrophysiological signals measured using electroencephalography (EEG) [10–12] and magnetoencephalography (MEG) [13].

Electrophysiological signals accessible from the scalp are particularly promising targets for non-invasive stimulation, as an alternative therapy to deep brain stimulation (DBS) of the ventrolateral thalamus. DBS provides an adaptable and bilateral therapy for ET, yet by its invasive nature, is only available to the most severely affected patients. To date, non-invasive brain stimulation has targeted the primary motor cortex [14,15] or the cerebellum [16–19], with the latter demonstrating superior clinical outcomes. Despite this, most approaches to stimulation remain open-loop and require high stimulation energies that can be uncomfortable for the patient. The development of next generation neurostimulation for ET requires (a) targets that can be accessed with minimally invasive techniques, and (b) biomarkers that can provide closed-loop, on-demand stimulation proportional to symptom severity.

The substantial overlap of the tremor circuit with volitional motor control networks [20,21] implies that common signals, such as 14-30 Hz beta band oscillations, could predict tremor state. For example, movement associated event related desynchronization (ERD) [22] of the beta band has been used to trigger DBS in tremor [23–25] at the onset of action or postural tremors, offering a method for anticipatory control. However, it remains unclear how the aberrant synchronization at tremor frequencies changes these physiological markers of motor processing. Previous studies have demonstrated that beta ERD over the primary motor cortex is significantly increased during movement in patients with ET or Parkinson’s disease using electrocorticography [26,27]. The narrow spatial coverage of these surgical recordings cannot address whether the observed changes align with the broader cortical disruptions reported in functional MRI studies [28,29]. It is also unknown whether changes are pathological or compensatory [30]. Furthermore, most studies to date have adopted a static paradigm, focusing on tremor during unchanging postures, and thus ignoring the dynamics of motor control in ET, especially during active limb movements such as reaching [31–33].

Motivated by a need to identify new biomarkers of tremor state and movement in ET, we hypothesised that the pathophysiology of ET disrupts the large-scale motor dynamics associated with naturalistic movement, such as reaching. To investigate this, we performed high resolution neuroimaging with high-density EEG and, validated for the first time in movement disorders patients, optically pumped magnetoencephalography (OPM) [34]. We first aimed to identify overlapping neural circuits activated by both whole limb reaching movements and pathological tremors. To aid interpretation of high-resolution neuroimaging recorded across multiple regions and frequencies, we use time frequency principal component analysis (tfPCA) [35]. This dimensionality reduction method allows us to decompose time locked, movement responsive oscillations recorded over the brain, into simplified components with associated *latent* dynamics. These low-dimensional representations capture how the brain coordinates during movement and tremor, linking regions and frequency bands into ensemble circuits with correlated activities. This can be thought of as analogous to identifying the separate sections (e.g. brass, strings etc) of an orchestra working together. Using these tools, we tested our central hypothesis: that changes in cortical networks in ET alter the coordination of large-scale oscillatory networks during movement.

## 2 Results

### 2.1 Behavioural Analyses of the Delayed Reach-to-Target Task

We recruited patients diagnosed with ET according to the clinical MDS-criteria [36] and age matched controls for recordings with either 128 channel EEG or a whole head OPM array. All patients displayed a mild to moderate upper limb tremor, scoring between 3 and 6 on “The Essential Tremor Rating Assessment Scale” (TETRAS, Elble, 2016). Clinical details and relevant medication are provided in tables in Supplementary Information I and II.

All participants performed an upper limb reaching task outlined in Figure 1A. Four of 31 EEG participants were excluded as they did not follow the task sequence, and four additional EEG participants were removed due to large muscle/movement artefact. This left a total of 23 subjects for analysis (11 healthy controls, 12 ET patients). Nine additional participants were recorded with OPMs (5 controls, 4 ET patients) with none excluded.

**Figure 1.**
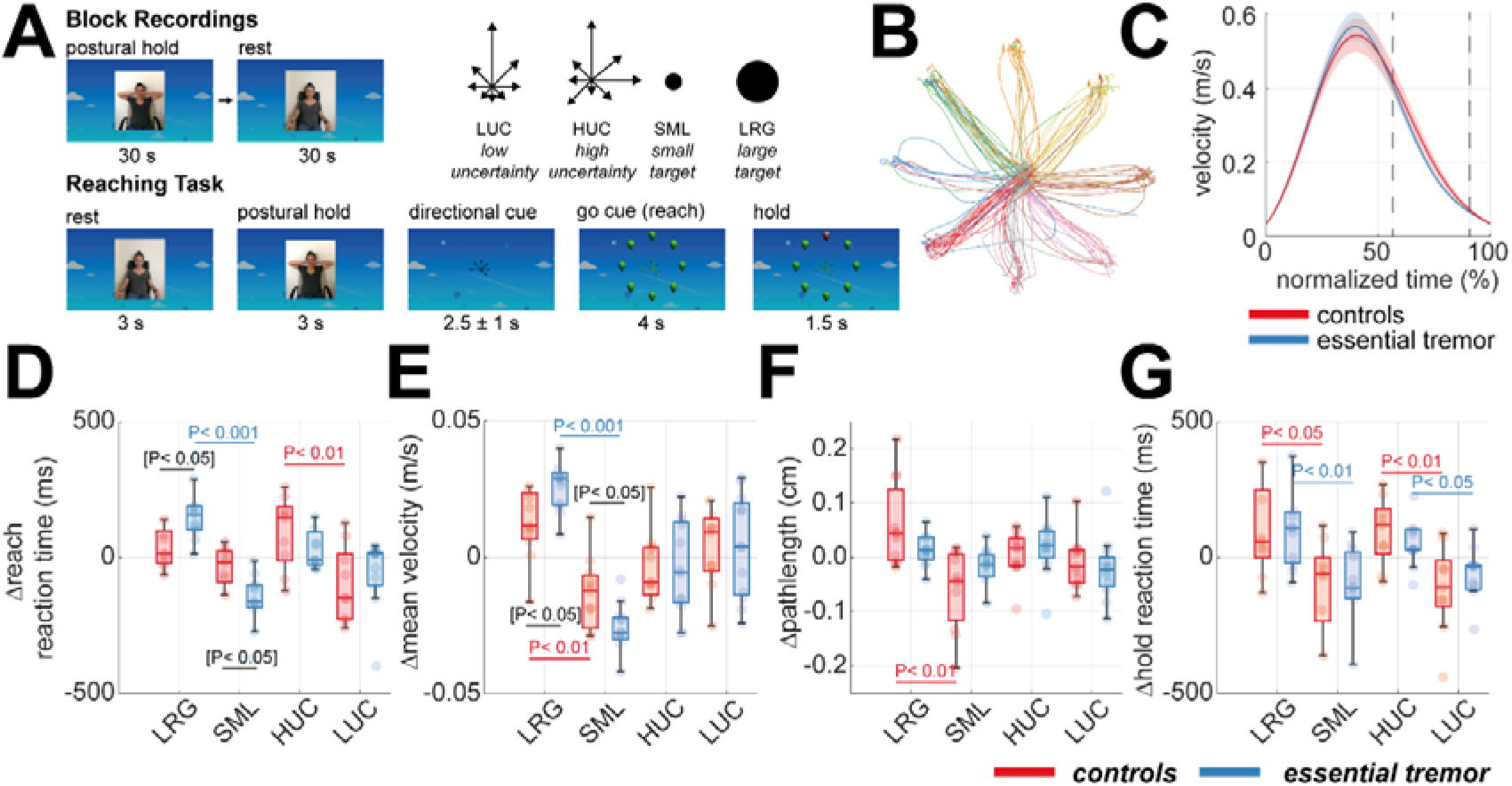
–Structure of delayed reach-to-target task and kinematic analysis. **(A)** Illustration of the sequence and timings of the reaching task. Each block began with 30 s of postural hold and eyes open rest. The reaching task consisted of *rest* (3 s), *posture* (3 s), onset of a directional *cue* (2.5 ±1 s), reach *execution* (max. 4 s), and sustained *hold* (1.5 s). The task followed a 2x2 design: high versus low uncertainty (HUC vs LUC), and large versus small targets (LRG vs SML). **(B)** An example set of trajectories from a single subject for the centre-out reaches, colour coded by the target. **(C)** The group averaged velocity profiles of the reaches, plot in normalized time from reach onset (0%) to hold onset (100%). **(D- G)** Boxplots summarising the between-subject statistics of task kinematics. Each point gives the deviation from the subject’s mean value. Data is shown for EEG experiment controls (red) and ET patients (blue), with colour matched bars indicating significant post-hoc t-tests. Black bars demark significant post-hoc t-test when comparing ET and controls. Brackets indicate tests that did not survive Bonferonni correction for multiple comparisons. Data shown for the EEG cohort only.

ET and control subjects’ reach kinematics were similar, with comparable reaction times and path lengths. ET patients reached with slower average velocities (ANOVA (84), F-statistic = 4.76, P < 0.05). To focus on within-subject variance, we subtracted the subject mean before computing task-related differences. Smaller target sizes resulted in slower reaction times in both groups (Figure 1D; ANOVA (82), F-statistic = 37.7, P < 0.001). Slower reach velocities were observed with both high cue uncertainty and smaller targets (Figure 1E; Uncertainty: ANOVA (84), F-statistic = 14.1, P < 0.001; Size: ANOVA (84), F-statistic = 7.39, P < 0.01). Only control participants showed changes in path length with target size (Figure 1F; t-test (20), t-statistic = 3.66, P < 0.01), indicating trajectory adaptation for small targets.

These results show that high cue uncertainty and high target precision modulate all subjects’ reaching kinematics, with ET patients making slower movements overall and showing more variation in velocity in response to changes in required motor precision, relative to controls. We return to this in Section 2.7 where we analyse how variations in oscillatory network dynamics can explain these behavioural variations.

### 2.2 Analysis of Upper Limb Tremor

Upper-limb tremor amplitude was estimated from accelerometery (Supplementary Figure 1). ET patients on average exhibited a 0.85 ± 0.98 m/s^2^ increase in tremor amplitude from rest to postural raise (paired t-test (10); t-statistic = 4.69; P < 0.001) at an average frequency of 5.8 ± 1.6 Hz. In 4/12 patients, a small (> 0.1 m/s^2^) rest tremor was also present. Control subjects displayed a small physiological tremor at posture of ∼0.19 ± 0.04 m/s^2^ (paired t-test (10); t-statistic = 7.06; P < 0.001). For reference, thresholds for visual detection of tremor are ∼0.07m/s^2^ [38].

Postural tremor amplitudes positively correlated with clinical scoring (TETRAS performance subscale; Spearman’s (12); R = 0.75; P = 0.010). There was no significant effect of cue uncertainty or target size on the tremor amplitude as evaluated at either motor preparation or the hold period (Supplementary Figure 2).

### 2.3 Peripheral Tremor Rhythms are Synchronized to Motor Associated Brain Regions

Our first analysis localized sources in the brain synchronized to tremor using a Dynamic Imaging of Coherent Sources (DICS) beamformer. In 8/12 ET patients measured with EEG, a peak in coherence (4-12 Hz, >85th percentile of the whole brain; binomial test, P < 0.001) was found over the cSMA (Figure 2A and B, EEG and OPM, respectively). This peak was also significantly higher in ET patients compared to controls (Figure 2C; peak t-statistic (10) = 2.24). A cSMA peak was also seen in OPM patients (Figure 2D; peak t-statistic (3) = 3.50). In a subset of patients, tremor coherence was found in the contralateral dlPFC (5/12 EEG cohort, binomial P < 0.05; 2/4 OPM cohort, binomial P > 0.05), that was found to be significantly greater in ET patients compared to controls (EEG cohort only, Figure 2C). In the OPM cohort, 3/4 patients had a tremor source in the contralateral PPC (binomial P < 0.05), that was also significantly elevated in ET (Figure 2D).

**Figure 2.**
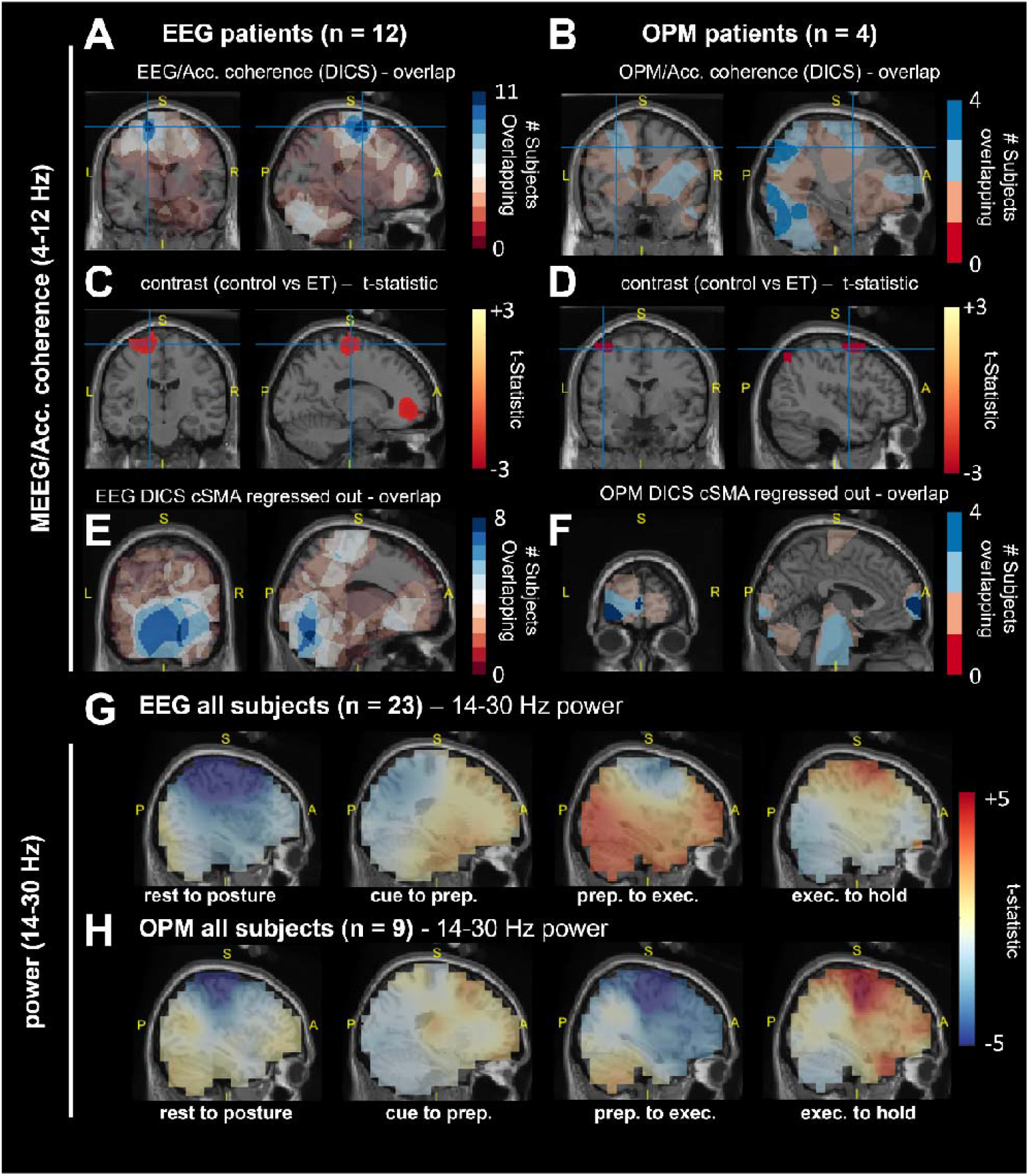
– Source maps of OPM/EEG data coherence with the 4-12 Hz peripheral tremor signal estimated using a Dynamic Imaging of Coherent Sources (DICS) beamformer from postural hold data and movement related beta (14-30 Hz) power changes. Subjects’ anatomy was flipped such that all slices were aligned contralateral to the dominant hand prior to performing statistics. **(A and B)** EEG and OPM image of the overlap of the top 90^th^ percentiles subject level coherence at source level. The slice location corresponds to the peak of the ET group average peak at the contralateral supplementary motor area (cSMA). **(C and D)** T-contrast of EEG and OPM DICS images comparing coherence in tremor band between control and ET. Maps were thresholded at the critical T-value (P < 0.05, uncorrected). Positive T-statistics indicate control > ET. **(E and F)** Same as (C and D) but for overlaps computed from the auxiliary DICS analysis, performed after regressing out the cSMA virtual channel. **(G and H)** Grand averaged EEG and OPM source power images across both controls and ET patients. Contrasts show unthresholded t-statistics using a baseline period at each stage of the reaching task.

An auxiliary DICS analysis, designed to detect weaker sources by regressing out the cSMA signal and focusing on the top 50% of coherent trials, revealed a tremor-coherent source in the ipsilateral and contralateral cerebellar lobule VI in 8/12 EEG patients (Figure 2E; binomial P < 0.001), that was significantly increased in ET vs controls (peak t-statistic (21) = 2.43). Additionally, a dlPFC peak was found in 4/4 OPM patients (Figure 2F; binomial P < 0.001). Sources varied across patients, with coherence peaks found in the cerebellum VI, cPPC, or dlPFC. There were no significant differences in kinematics when ET patients were divided by these peak regions (ANOVA (11); P > 0.05). The cerebellar group tended to have more severe tremor although this did not pass significance (t-test (8), t-statistic = 1.27; P > 0.05). These results identify ROIs in the cSMA, cPPC, dlPFC and Cerebellum VI that are synchronized to tremor in ET patients and are consistent with previous reports [10,11,13]. Importantly, these brain areas also exhibit significant movement associated activity in the beta frequency (Figure 2G and H; Supplementary Figure 3) suggesting the potential for ET pathophysiology to influence healthy motor oscillations. Most notably, our time frequency analyses show that cPPC exhibits a clear desynchronization during a period of motor planning following cue onset that is apparent in both controls and patients (Supplementary Figure 4 and 5) In later analyses, we will use dimensionality reduction to capture the collective dynamics of induced oscillations across motor networks including composed of the regions identified with DICS here (Section 2.5).

### 2.4 Essential Tremor Patients Exhibit Deeper Movement Related Beta Desynchronization During Reaching

To investigate whether ET pathophysiology impacts physiological signals associated with motor processing, we first focused on the activity in the SMA, as this region was synchronized to tremor activity and exhibited motor induced responses in the majority of subjects recorded (Section 2.3). Relative to controls, ET patients show a 15% increase (relative to at rest) in lower beta frequency (14- 21 Hz) ERD in SMA (Figure 3A) during reach execution (two-sided t-test, permutation statistic (21), P* = 0.021) and the post-movement hold period (two-sided t-test, permutation statistic (21), P = 0.003).

**Figure 3.**
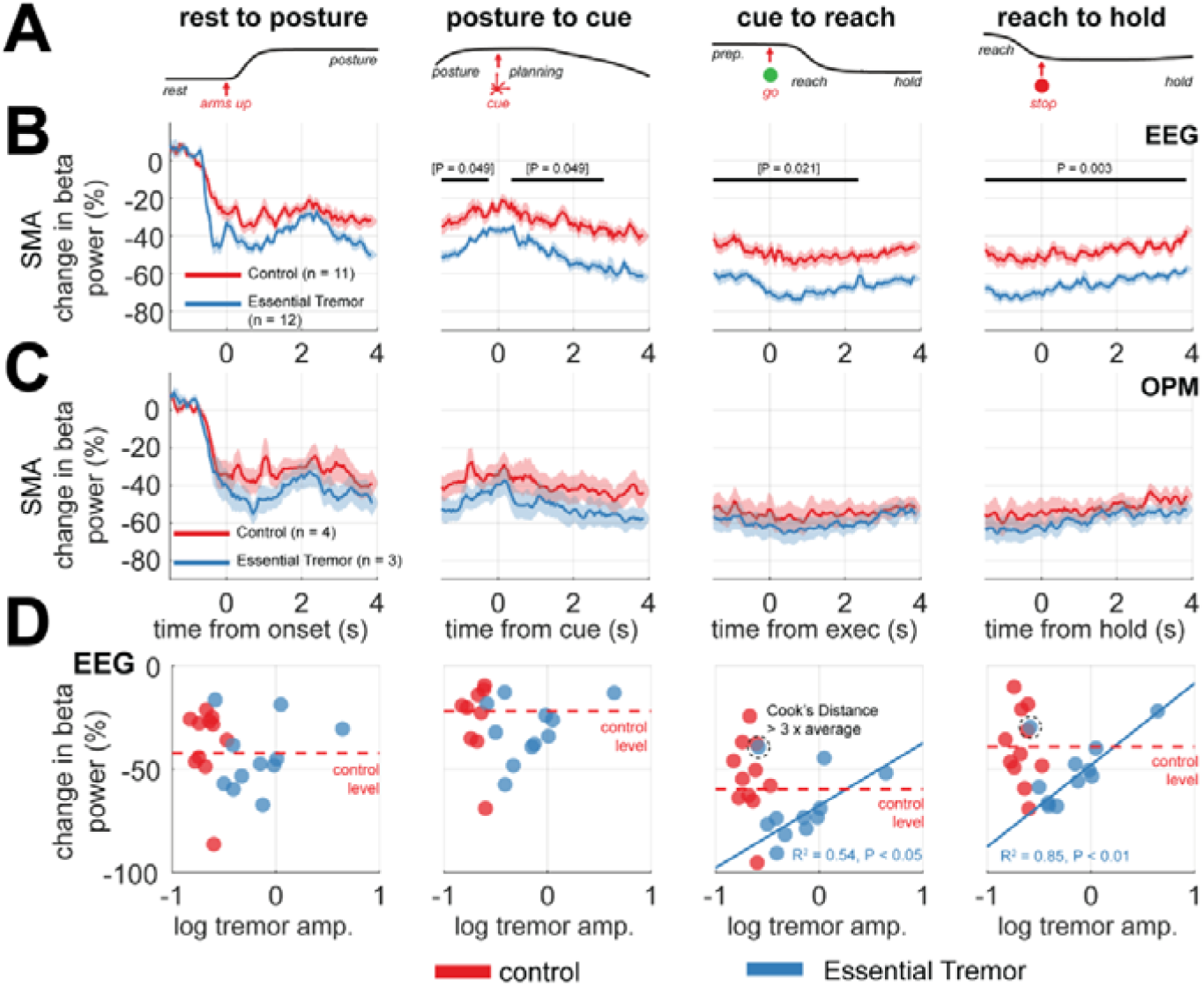
– **Analysis of movement locked beta event related desynchronization (ERD) in the supplementary motor area (SMA) comparing between Essential Tremor (ET) patients and control recordings in EEG. (A)** The grand- averaged accelerometer trace indicates the sequence of the task. **(B)** Time courses of low beta power (14-21 Hz) of the combined left and right SMA, scaled as the percentage change relative to at rest (0%), shown across the postural hold, reach planning, execution, and the hold period (columns left to right). Data is shown as the group average ±SEM, for controls (red) or ET patients (blue). Bars indicate significant decrease between controls and ETs, determined by cluster permutation statistics (P < 0.05). Brackets indicating tests that did not survive Benjamini and Hochberg False Discovery Rate correction for multiple comparisons. **(C)** Same as (A), but for OPM data. **(D)** Scatter plots of log scaled tremor amplitude versus the percentage change in beta power. In the case of significant Pearson correlation, bold blue lines indicate the associated regression for ET patients only. The circled point indicates the outlier (identified according to Cook’s criteria) removed from regression.

ET patients studied with OPM also exhibited greater movement-related low beta frequency ERD to controls (Figure 3B), although this did not reach statistical significance, likely due to the small sample size. The degree of SMA beta desynchronization during reach execution and the post-movement hold period were both negatively correlated with overall subject averaged tremor severity (as computed from 45s blocks of continuous postural hold) of ET patients in the EEG cohort (Figure 3C; reaching: R^2^ = 0.54, P < 0.05 and hold: R^2^ = 0.85, P < 0.01; one outlier removed on Cook’s distance). These findings support our hypothesis that ET pathophysiology disrupts physiological neural activity such as that found in the SMA. In the following sections, we examine whether this phenomenon is localized or extends to a broader motor network.

### 2.5 Time-Frequency Principal Component Analysis Identifies Known Motor Circuits

To understand how ET may affect large scale motor induced oscillations, we used time-frequency PCA (tfPCA) [35] to summarise a large set of time-frequency descriptions of M/EEG source activity estimated across regions of the brain identified to synchronize to tremor and exhibit motor responsive beta oscillations.

When applied to the group averaged EEG data (all ET and control subjects), tfPCA yielded four components with distinct spatial and spectral weights (Figure 4B) that explained 97% of the total variance, and accurately reconstructed the original data (Figure 4H). These four components can be visualized in the original time-frequency space via back-projection (Figure 4C-H) or represented via their latent time dynamics (shown in Figure 5). Component (1) represents a fronto-parietal- sensorimotor network active in the lower beta band (14-21 Hz) and desynchronises at the onset of the movement cues (i.e. motor planning; 64% of total variance); Component (2) represents a sensorimotor network that exhibits movement locked high frequency gamma activity (> 30 Hz; 17% of the total variance); Component (3) represents a premotor/dorsolateral prefrontal network active at mu/beta band (8-16 Hz) that desynchronises at the onset of movement (i.e. motor execution; 10% of total variance); and Component (4) represents a premotor/dorsolateral prefrontal network active across the upper beta band (18-30 Hz) that desynchronizes at movement onset (i.e. motor execution; 6% of the total variance). Notably, our analyses unveil component dynamics that separate between movement planning (component 1) and preparation (component 2).

**Figure 4.**
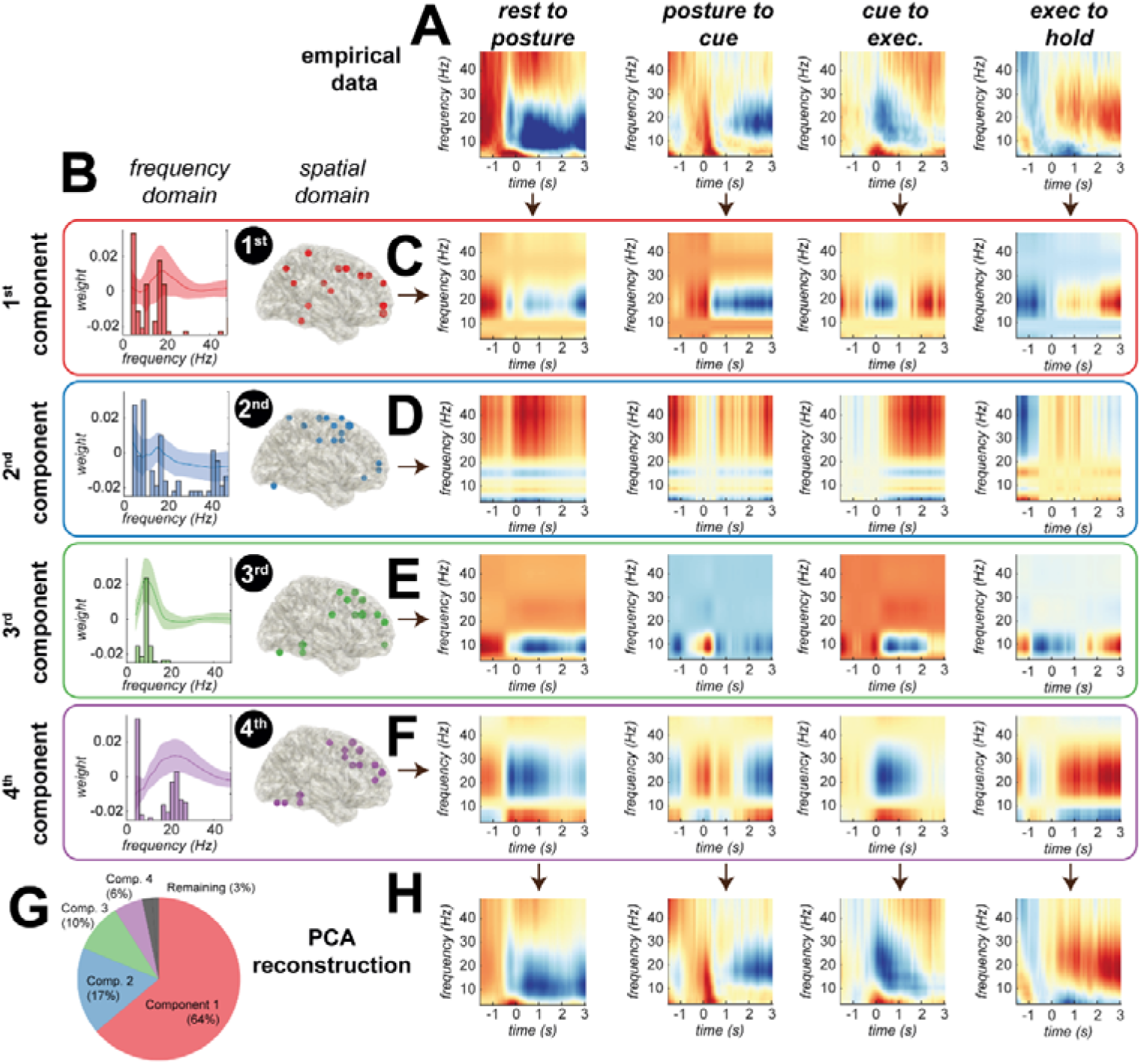
– Identification of large-scale, movement related brain circuits using time-frequency principal component analysis (tfPCA) applied to group averaged data (both ET and controls) derived from source-space projected EEG virtual electrodes. **(A)** Time-frequency spectrograms were computed for each region of interest (ROI) in the network, incorporating the dorsal prefrontal cortex, primary and supplementary motor areas, posterior parietal cortex, and cerebellum VI. tfPCA was applied to the group averaged EEG data, with spectrograms concatenated for each motor epoch. Prior to tfPCA, data was log scaled, and Z-normalized per subject. Visualization of the coefficients of each PCA component 1-4 (red, blue, green, and purple, respectively) in both the **(B)** frequency and spatial domains. **(C-F)** Back projection of empirical data allows for visualisation of the time-frequency dynamics of each component (averaged in space), across the four different epochs of the reaching task (corresponding to the rows). **(G)** Explained variance of the data by the four components. **(H)** Reconstruction of the original time-frequency data using the four components.

**Figure 5.**
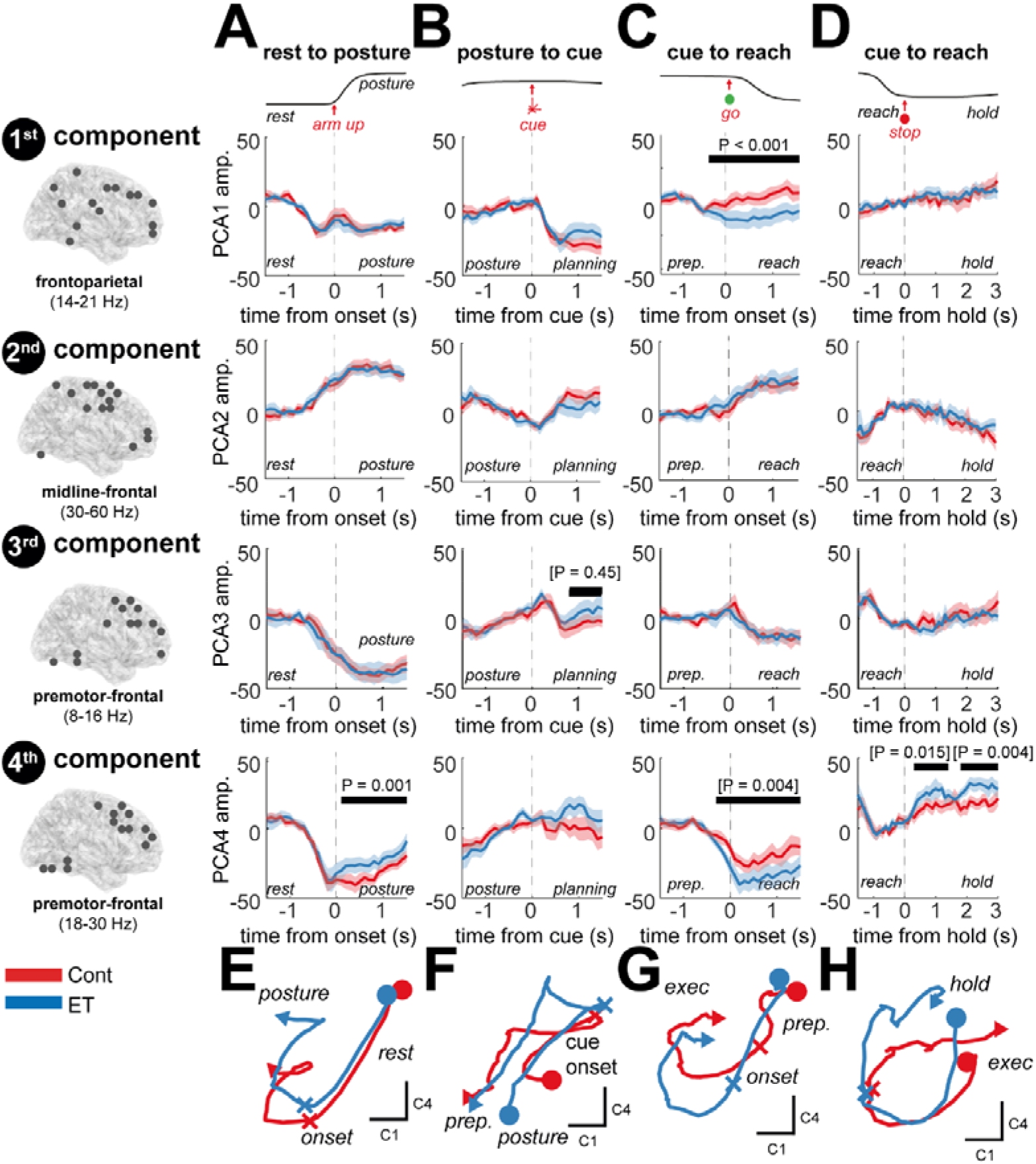
– Visualization of motor responsive, latent circuit dynamics and a comparison between controls and ET patients recorded with EEG. Components were computed using tfPCA applied to the group averaged EEG data (Figure 4). These coefficients were then used to project data to trial-level latent dynamics that could be used to explore differences between controls and ET patients. The grand averaged accelerometer traces are shown at the top to indicate the task sequence. **(A)** The latent dynamics exhibited during the postural hold for each component (columns) are plot for ET (blue) and controls (red) separately. Bars and associated P-values indicate the outcome of cluster permutation statistics between the two experimental groups. Brackets indicate tests that did not survive Benjamini and Hochberg False Discovery Rate correction for multiple comparisons (16 tests in total). **(B-D)** Same as (A) but for cue presentation, reach execution, and the sustained hold. **(E-H)** Plots of the 2D latent trajectories (components 1 and 4, only) indicate highly correlated dynamics between controls and ETs with quantitative differences in the weighting of the components such as increased engagement of the prefrontal beta network in ET subjects (4^th^ component, apparent in plot G). An equivalent analysis was performed for OPM recordings (Supplementary Figure 6).

### 2.6 Essential Tremor Patients Exhibit Differences in Large-Scale Circuit Dynamics

Using the tfPCA loadings computed at the group level we reconstructed trial-averaged latent dynamics for each subject (Figure 5A) and examined differences between cohorts (ET and age-matched controls). Latent dynamics were preserved between the controls and ET subjects (Figure 5E), with average between subject Pearson’s correlation coefficients of R = 0.91 and R = 0.84 for EEG and OPM, respectively (Supplementary Information III, table 1). Across EEG and OPM modalities, we found an average correlation coefficient of R = 0.73 (Supplementary Information III, table 2).

Our results show that in the EEG data, the frontoparietal low beta (14-21 Hz) network (1^st^ component) exhibits a cue locked desynchronization that does not differ between controls and ET patients (Figure 1A) in the planning stage (i.e., at cue onset). However, during movement execution, ET participants exhibit further desynchronization compared to controls (Figure 5C 1^st^ row; permutation t-statistic (74), P < 0.001). Both OPM and EEG dynamics exhibited an ERD at upper beta (18-30 Hz) frequencies in the dorsolateral prefrontal beta network (4^th^ component) that was increased in ET relative to controls, during execution of both the postural hold (Figure 5A 4^th^ row; EEG permutation t-statistic (74), P* = 0.001) as well as the reach (Figure 5C 4^th^ row; EEG permutation t-statistic (74), P* = 0.004). This was followed by a faster and stronger rebound during the post-movement hold period (Figure 5D 4^th^ row; EEG permutation t-statistic (74), P* = 0.004). These latent dynamics computed at the single subject level expand on ET associated increases in beta ERD in the SMA (Section 3.4) by highlighting that observed changes are associated with increased frontoparietal beta ERD following movement. This supports our principal hypothesis that ET results in altered latent dynamics of large-scale motor networks during reaching.

### 2.7 Whole Brain Latent Networks Can Explain Within-Subject Variation in Tremor and Kinematics

To understand how pathophysiological changes in motor dynamics may relate to functional processes, we analysed the association between latent dynamics and kinematic variables. To this end, we projected latent dynamics to the single trial level, using the same coefficients as before (Section 2.6), and made a statistical comparison between data split into *low* and *high* trials of each kinematic (1^st^ and 4^th^ quartiles, respectively). For statistical power, we looked only at the EEG cohort due to larger sample size. We analysed both the tremor power (in ET patients only), and the averaged reach velocity/movement variability (for combined ET and controls).

In Figure 6, we examine the frontoparietal and premotor networks (components 1 and 4, respectively), as these were most impacted in ET (Figure 5). This analysis shows that prolonged desynchronization of the low beta frequency (14-21 Hz) frontoparietal network during reaching (Figure 6C), is associated with reaches of slower average velocity (1^st^ component; green bar; permutation t-statistic (21), P = 0.005). This was not found to be related to the total duration of the reach. Furthermore, faster resynchronization of this same network (Figure 6D) increased the stability of movement in the hold period (red bar; permutation t-statistic (21), P = 0.007). These findings may help explain why the ET subjects exhibit overall slower reaches, when compared to controls.

**Figure 6.**
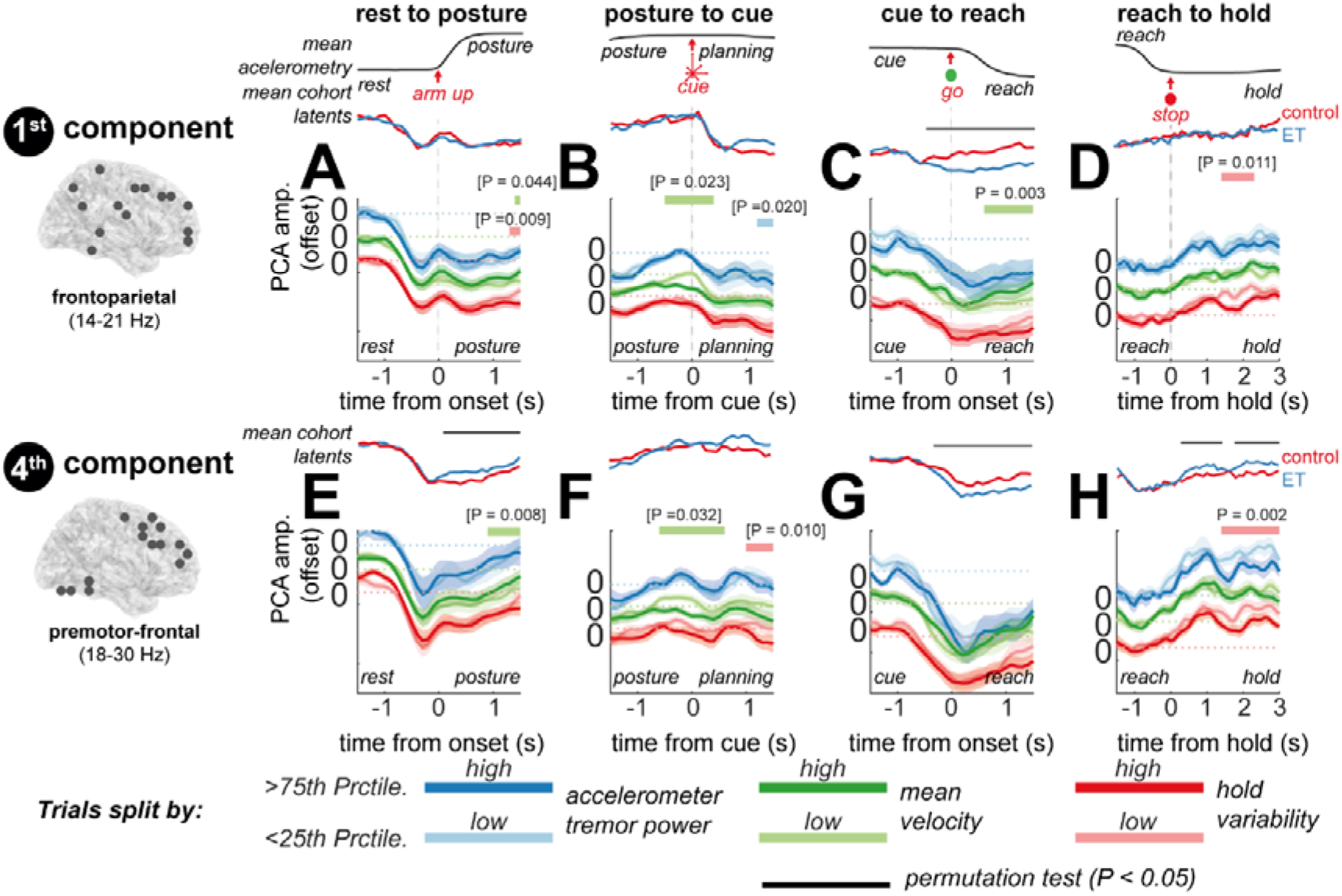
– Analysis of modulation of latent circuit dynamics altered in Essential Tremor, with movement kinematics during the reaching task. Latent states were projected to the single trial level, using coefficients computed from group averaged time frequency EEG data. Mean latent dynamics (reproduced from Figure 5) for controls (red) and ET patients (blue). Differences in average velocity/hold variability are shown for both controls and ET patients, whereas tremor parameters for ET patients only. Trials were split into 1^st^ (light colours) and 4^th^ quartiles (dark colours) and then subject averages compared at the group level using a cluster permutation test (indicated by thick lines above, and relevant P value for the test statistic). Brackets indicate tests that did not survive Benjamini and Hochberg False Discovery Rate correction for multiple comparisons (16 tests per parameter). **(A-D)** Plots of the latent dynamics of the 1^st^ component over the postural raise, cue onset, reaching, and hold, split by the tremor band power during each hold period (blue lines), the mean velocity during the postural raise/reach (green lines), and the hold variability (red lines). **(E-H)** Same as (A) but for the 4^th^ component. The 1^st^ and 4^th^ components are shown as these were most modulated by ET, the remaining components are presented in Supplementary Figure 7.

In the premotor-frontal network, faster resynchronization of upper beta frequencies (18-30 Hz) after reaching (Figure 6H) – a marker of ET as shown in the previous analysis – was associated with increased kinematic stability during the hold period (red bar; permutation t-statistic (21), P < 0.001). A similar effect was found during the postural hold period (red bar; permutation t-statistic (21), P* = 0.008) suggesting that premotor network can act to stabilize movement during static contraction. An increase in frontoparietal engagement during reaching suggests that ET subjects may increase their reliance on sensory circuits for online control during movement, a point we will return to in the discussion.

## 3 Discussion

Our results demonstrate that pathophysiological changes associated with ET lead to alterations of oscillatory dynamics within large-scale motor circuits during reaching movements. While ET patients executed movements with an accuracy comparable to controls, their reaching velocities were slower. Key motor regions were synchronized to tremor rhythms while also exhibiting movement-locked beta oscillations. Notably, the SMA - a region found to be highly synchronized to peripheral tremor – exhibited a 15% increase in beta desynchronization in ET patients that also inversely correlated with tremor severity. Whole-brain analyses using tfPCA revealed broader disruptions to a sequential pattern of latent network synchronization, particularly in frontoparietal and premotor networks. These networks, typically involved in visuomotor control and planning, showed enhanced beta desynchronization during movement in ET, which may reflect adaptive changes in patients that lead to slower reaching movements.

Our findings build on previous work linking SMA, PPC, and cerebellum to the tremor network [10–12]. These regions are also strongly coupled with ventrolateral thalamus at beta frequencies [39], suggesting the capacity of DBS to alter these networks. Cortically sensed signal offer the potential for closed-loop control of DBS, as demonstrated in Parkinson’s disease[40]. This approach could deliver thalamic stimulation tailored to expected tremor levels while accounting for motor context, enabling state-dependent neuromodulation in ET [41,42]. Regions such as the PPC and cerebellum also emerge as promising targets for non-invasive stimulation, consistent with recent therapeutic advances with scalp level electrical stimulation [18,19].

Beyond single-region analyses, our investigation of latent dynamics reveals the highly distributed nature of network disruptions in ET. These findings support the potential for multi-site sensing in closed-loop neuromodulation [43,44]. Furthermore, we demonstrate the strength of dimensionality reduction approaches, such as tfPCA, in improving the estimation of both motor and pathological states [45,46] via the identification of global network states [47]. Supervised methods for dimensionality reduction are likely to improve this further by focusing latent dynamics on context relevant signals [48,49]. Using these simplified, low dimensional representations to predict tremor severity, could enable proportional controllers for DBS, as has been demonstrated in Parkinson’s disease[50].

Movement responsive neural oscillations in the healthy brain [22,51,52] reflect important elements of sensorimotor processing [53]. Our finding of increased beta ERD in ET suggests that the brain adapts to mitigate the effects of hypersynchronous tremor rhythms. Increased beta ERD may suggest that tremor engages mechanisms similar to intentional movement, and/or reflect an adaptation that allows cortical neurons to encode movement in the face of tremor-related entertainment. By freeing up cortical neurons to encode vital parameters of movement[54], additional ERD may compensate for extraneous entrainment at the tremor rhythm (illustrated in Figure 7). Notably, differences in beta ERD were most pronounced during goal-directed reaching, aligning with evidence that more complex tasks demand greater cortical resources [55]. In severe tremor, synchronization to tremor may saturate the neuronal pool, leaving fewer neurons available to desynchronize at beta frequencies, which may explain the weaker motor-related ERD observed in the more severely affected OPM cohort. If neuronal dynamics adapt locally, it is crucial to understand how these changes propagate to produce the altered global oscillatory dynamics observed in this study. Recent research demonstrates that low- dimensional representations of neural population activity—vital for movement control[56] and correlated with oscillatory neural dynamics [57]—that are likely to influence distributed effects via long-distance synchronization.

**Figure 7.**
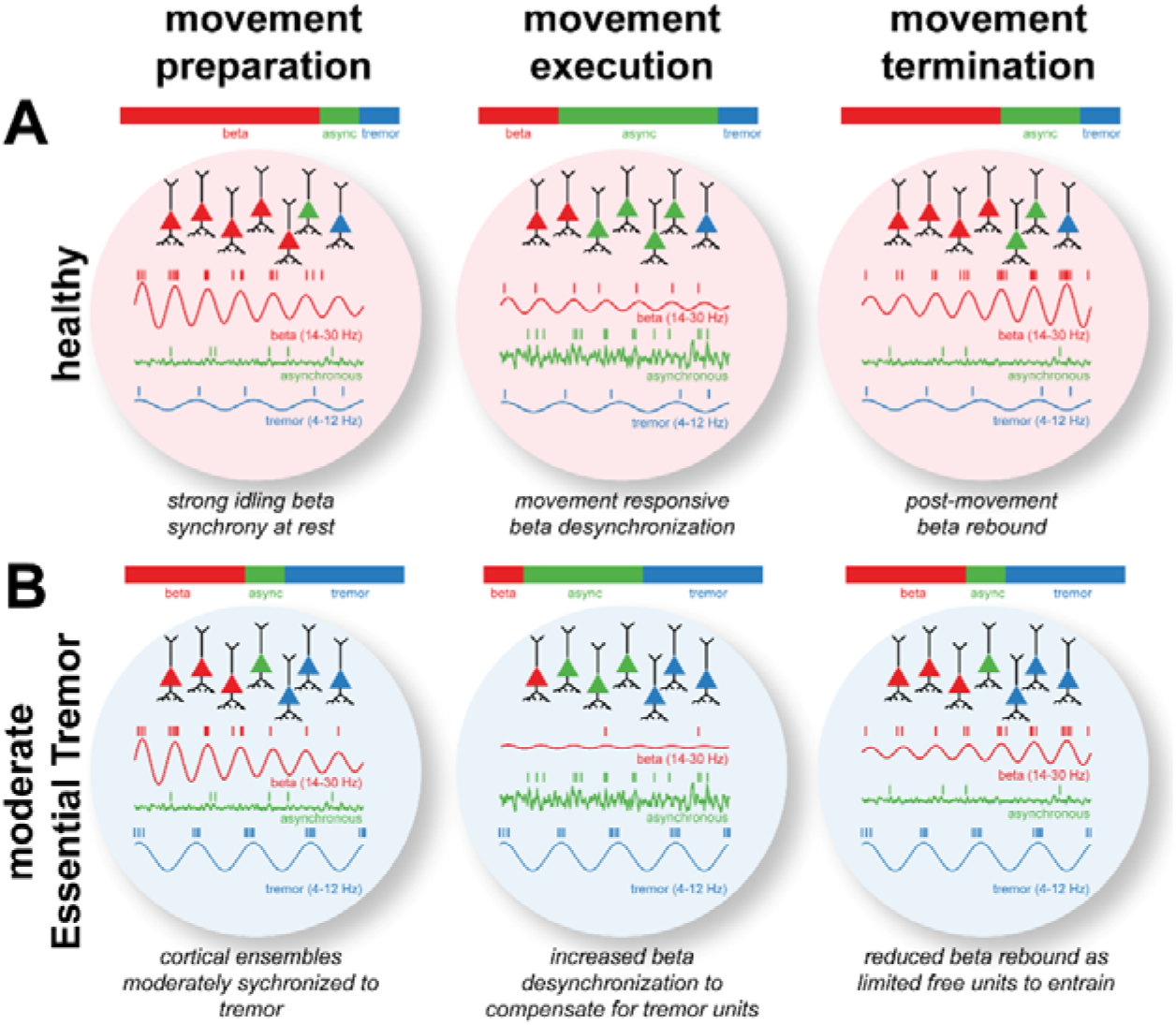
– Hypothetical neuronal mechanisms underlying alterations to motor induced dynamics following pathological synchronization in Essential Tremor. (A) During movement preparation (first column), neurons entrain to brain rhythms such as the sensorimotor beta rhythm (shown in red), but also tremor (blue). During movement (middle column), motor cortical neurons exhibit firing rate changes (assumed to be asynchronous; shown in green) that encode movement parameters. Simultaneously, ERD of the beta rhythm relinquishes units to participate in motor encoding, followed by a post movement beta rebound (final column). (B) In moderate ET, the tremor rhythm entrains a fraction of cortical neurons reducing the effective information encoding capacity of the available pool. To compensate, during movement, increased beta ERD frees up the neuronal “real estate” required to effectively encode movement.

Beta synchrony in the sensorimotor system is thought to reflect sensory gating [58–60], and post-movement error updating [61,62]. In ET, it is likely that cerebellar deficits introduce a susceptibility to sensorimotor noise [63,64]. We hypothesise that enhanced post-movement rebound could reflect compensatory suppression of sensory information. Notably, we found that the frontoparietal circuit exhibited an additional desynchronization in ET, a network that is most typically associated with visuomotor control [65]. These changes may explain the overall decrease in average movement velocity in ET patients. We hypothesize that this reflects a disruption of feedback control in ET patients. This accords with findings that online control of movement is impaired [66], while coupling between the parietal and motor cortices is disrupted in ET[29]. Our observations of effects across large-scale networks raise the question of whether DBS for ET can restore altered movement processing, as has been explored in the context of Parkinson’s disease [67]. Previous modelling work has shown that patterning of stimulation can be tuned to achieve specific network states [41], suggesting the potential for adaptive stimulation to steer adaptive motor activity in ET.

This work has investigated whole limb naturalistic movements and faces the well-recognised challenges of motion-related artefacts arising from either contamination by myographic activity and cable noise. Measurement of tremor during large movements is challenging, due to varying tremor axis and gravity artefacts, that could be avoided using gyroscope estimates of tremor severity in future studies. Additionally, there was no explicit forward model of the cerebellum, a component that would likely improve the accuracy of cerebellar localization in future studies [68].

This study reveals large-scale motor network disruptions in ET, with increased beta ERD across the circuit acting to compensate for neural resources entrained at tremor frequencies. For the first time, we have shown that frontoparietal beta rhythms are altered by ET pathophysiology and reflect increased slowing of reaching movements that may be explained by an increased dependence on feedback control. Our findings have broader implications for understanding not only ET but also adaptive mechanisms in the motor system associated with other pathologies, such as Parkinson’s disease. These results provide novel regions and biomarkers that can inform the next generation of brain stimulation therapies for tremor.

## 4 Methods

### 4.1 Recruitment and Details of Participants

Ethical approvals were obtained from the committees of the Philipps-University Marburg and University College London. Consent was obtained according to the Declaration of Helsinki. EEG experiments were conducted with 16 patients (51 ±19 years of age, 7 female) diagnosed with essential tremor (ET) [36] and 15 controls (42 ±16 years of age, 9 female). OPM was conducted with 4 ET patients (50 ±20 years of age, 1 female) and 5 healthy controls (50 ±13 years of age, 1 female). The EEG cohort (Marburg) had a mean TETRAS performance score of 18.7 ±8.2 and the OPM cohort (UCL) had scores of 19.5 ±3.8. A subset of the EEG cohort displayed mild head tremor (5 of 12 included patients scoring 1 or 2).

### 4.2 Delayed Reach-to-Target Task

Participants performed a delayed reach to target task, making whole limb reaches to mimic naturalistic movement. The task (Figure 1A) required participants to adopt a 90-degree upper arm elevation against gravity with flexed elbow posture and then, from this position, make centre-out reaches to one of eight targets and hold their finger fixed to “pop” target balloons. The task sequence was: (a) 3 s eyes-open rest, (b) 3 s postural (as above) hold, (c) 2.5 ±1 s movement cue presentation, (d) appearance of a GO cue, (e) 1.5 s of sustained hold. A point was scored if the correct balloon was popped (as predicted from the dispersion of arrow cues). Further details of the task design can be found in Supplementary Information III.

### 4.3 High Density Electroencephalography

High-density EEG (HD-EEG) was recorded in a recording chamber using an elastic cap to mount active, gel-based electrodes (Brain Products GmBH, Gilching, Germany) in the standard 10-10 system with 128 electrodes and amplified using a Brain Products DC amplifier. Electrode gel was applied to maintain electrode impedance below 10 kΩ. All data were recorded with a 1 kHz low pass filter and digitized at 5 kHz. EEG recordings were made with a reference at FCz, which was re-referenced offline to the common average.

### 4.4 Optically Pumped Magnetometer Recordings

Optically pumped magnetoencephalography (OPM) was made using a combination of 2^nd^ and 3^rd^ generation QuSpin sensors (dual- and tri-axial sensors, respectively; QuSpin Inc., Louisville, Colorado, USA) mounted in rigid 3D-printed casts (Chalk Studios, London, UK) custom-built to each participant’s scalp shape determined from structural magnetic resonance images (MRIs). In the absence of an MRI (1 control and 1 patient), the head shape was estimated using an infrared depth camera. Offsets of 1-3 mm were added to scanner casts to allow for anatomical error, tissue deformation, and hair.

OPM experiments were conducted in a magnetically shielded room (MSR; Magnetic Shields Ltd, Staplehurst, UK). The inner layer of mu-metal lining the room was degaussed using a low-frequency decaying sinusoid driven through cables within the walls before the start of the experiment. The OPM sensors were then nulled using onboard nulling coils and calibrated. The OPM acquisition system (National Instruments, Texas, USA) had a sampling frequency of 6 KHz and a 16-bit resolution. An antialiasing 500 Hz low-pass filter (60th order FIR filter combined with a Kaiser window) was applied before data were down-sampled offline to 2 kHz.

### 4.5 Kinematics and Accelerometery Recordings

Arm movement was captured with Optitrack Motive software (Planar Systems, Oregon, USA) and an infrared camera (Optitrack Duo/Trio) tracking a reflective marker attached to the participant’s hand. Task triggers were sent to the EEG amplifier or the OPM DAC using a LabJack U3 DAC (LabJack Corp., Colorado, USA). Acceleration was recorded via a triaxial sensor (EEG, Brainproducts GmBH; OPM; ADXL335; Analog devices Inc., Wilmington, Massachusetts, USA) affixed to the affected limb. For one EEG patient, accelerometery was not available. Therefore, this participant is excluded from analyses requiring estimates of tremor (EEG/Acc. coherence and trial-by-trial tremor analyses). Statistics of movement kinematics were computed by averaging trials over each condition, removing outliers with |Z-score| > 1.96, and then computing an ANOVA.

### 4.6 Preprocessing, Artefact Removal, and Data Rejection

Analysis was performed in MATLAB (The Mathworks, Massachusetts, USA) using the Fieldtrip[69] and SPM 12 toolboxes. Digitized data were down sampled to 512 Hz. For EEG, individual channels were inspected to remove those affected by gross movement artefact, and then high and low pass filtered at 2 and 98 Hz using FIR windowed-sync filters. Line noise was removed by fitting sine and cosine components to the line frequency and then subtracting from the data. Oculomotor, muscle, or cable artefacts were removed using independent component analysis (ICA). Trials with remaining artefacts were rejected visually, using Z-score and kurtosis measures set at |Z| > 5; and |K| > 6. Bad channels were replaced via local spline interpolation. OPM signals were processed identically but used adaptive multipole model [70] for interference reduction following ICA.

Participants were removed when task sequence was not well followed, greater than 75% of reaches being rejected by the CNN, or EEG data quality being poor (high prevalence of cable, movement or muscle artefact).

### 4.7 Time Locking and Spectral Analysis of Data

The onsets of each postural raise were found from thresholding the Z-normalized sum of the triaxial accelerometers at Z = 3. Reach onsets and holds were marked using a semi supervised approach in which three independent scorers manually marked accelerometer and motion tracking traces, provided a quality score of 1-3 for each trial in a random subset of the data (30%). For precise criteria, please see Supplementary Information V. These labels were then used to train a convolutional neural network (CNN) that we deployed on the full dataset. Performance assessments of the CNN are given in Supplementary Figure 8. Time-frequency representations were calculated using Slepain multitapers, scaling the time window to 4 cycles per frequency bin, utilizing a 0.4 Hz smoothing window, and incorporating 10 seconds of zero padding.

### 4.8 Source Space Analysis

Dynamic Imaging of Coherent Sources [71] (DICS) mapped the distribution of power or coherence across the cortical volume. Wideband (2-98 Hz) beamformer weights were used to project sensor-level data to source-level virtual electrodes. To maximise sensitivity of DICS to weakly synchronized sources, an auxiliary analysis used trials from the top 50% of coherence for contralateral supplementary motor area (cSMA; the peak of the group averaged image), and then regressed out the cSMA virtual channel signal, following [10]. Brain parcellations used the Automated Anatomical Labelling atlas 3 [72]. For full details concerning the construction of forward models and sensor coregistration, please see Supplementary Information VI.

Statistics in the source space were computed using either one-sample (DICS coherence in tremor band), or two-sample (ET/Control contrasts, cue locked contrasts) t-statistics. DICS images were first normalized to account for unequal sample size [73,74]. Statistical maps are presented as (a) overlap maps of the top 85^th^ percentile of each subject; and (b) t-statistics thresholded at the critical level (P < 0.05, uncorrected. The probability associated with overlaps was derived from a binomial distribution B(k > n; N, p), indicating the probability of observing >*k* overlaps out of *N* subjects, where *p* represents the probability of overlap (*p* = 0.15, for thresholding at the 85^th^ percentile).Uncorrected source statistics formed an initial analysis to define regions of interest (ROIs) that we carried forward to test our main hypothesis regard time-frequency domain activity.

### 4.9 Estimation of Tremor Amplitude

Tremor signals from triaxial accelerometery (Supplementary Figure 1) were converted to units of acceleration (m/s^2^) using the manufacturer’s conversion factors. Tremor was measured at the following epochs: during the postural hold, post-cue onset, and after termination of the reach. For each epoch, we computed the principal component of the filtered triaxial accelerometer signal (4-12 Hz; bandpass FIR filter) [3]. The peak tremor frequency was then computed from the multitapered power spectrum (1.5 tapers/Hz) of the first component. An estimate of the tremor amplitude was made by computing the root mean square of the dominant axis of the accelerometer signal, after bandpass filtering around the tremor peak ± 3 Hz. Correlations were checked for outliers using three times the mean Cook’s distance as a threshold [75].

### 4.10 Identifying Latent Spectro-Spatial Components with Time-Frequency Principal Component Analysis

Large-scale brain circuits associated with voluntary movement control and tremor were identified using time-frequency Principal Components Analysis (tfPCA)[35]. tfPCA projects time-frequency data computed at voxels across the brain (*time* x *frequency* x *space* x *repetitions*) to a low dimensional set of components with associated latent dynamics (*component* x *time*). Voxels informed from the DICS source analysis included ROIs in the ipsi- and contra- lateral dorsal prefrontal cortex, primary and supplementary motor areas, posterior parietal cortex, and cerebellum lobule VI. tfPCA was applied to the group averaged motor induced responses with weights used to back-project data to the single trial level. Further details are given in Supplementary Information VII. To test changes associated with kinematics or tremor, we applied cluster-based permutation testing to data split by the 1^st^ and 4^th^ quartile of each subject, using P = 0.1 as a cluster forming threshold. This threshold is not the alpha level for cluster test significance (set at α = 0.05), but the threshold needed to carry a cluster forward for permutation testing.

## 5 Data Availability

Upon acceptance, the MATLAB code describing the analyses in this report will be shared to a publicly available version-controlled repository on Zenodo. The current repository is available at: https://github.com/twestWTCN/NetworksInEssentialTremor. The data for this study is not currently available for dissemination due to the terms of our clinical ethics, if passing editorial approval we can seek secondary ethical approval to obtain consent for further sharing of this data.

## Acknowledgements

We extend our thanks to the UCL OPM development team for making these recordings possible; Carolina Reis for help with early testing; the authors who develop, maintain, and freely share several toolboxes (listed in Supplementary Information VIII); Alto Neuroimaging at Aalto University for providing the accelerometer system used in the OPM experiments; and the anonymous participants who generously gave up their time to take part in this study.

