## Supplementary Information for "Essential Tremor Disrupts Rhythmic Brain Networks During Naturalistic Movement"

### 1 Supplementary Figures

#### 1.1 Supplementary Figure 1 – Analysis of Essential Tremor Properties

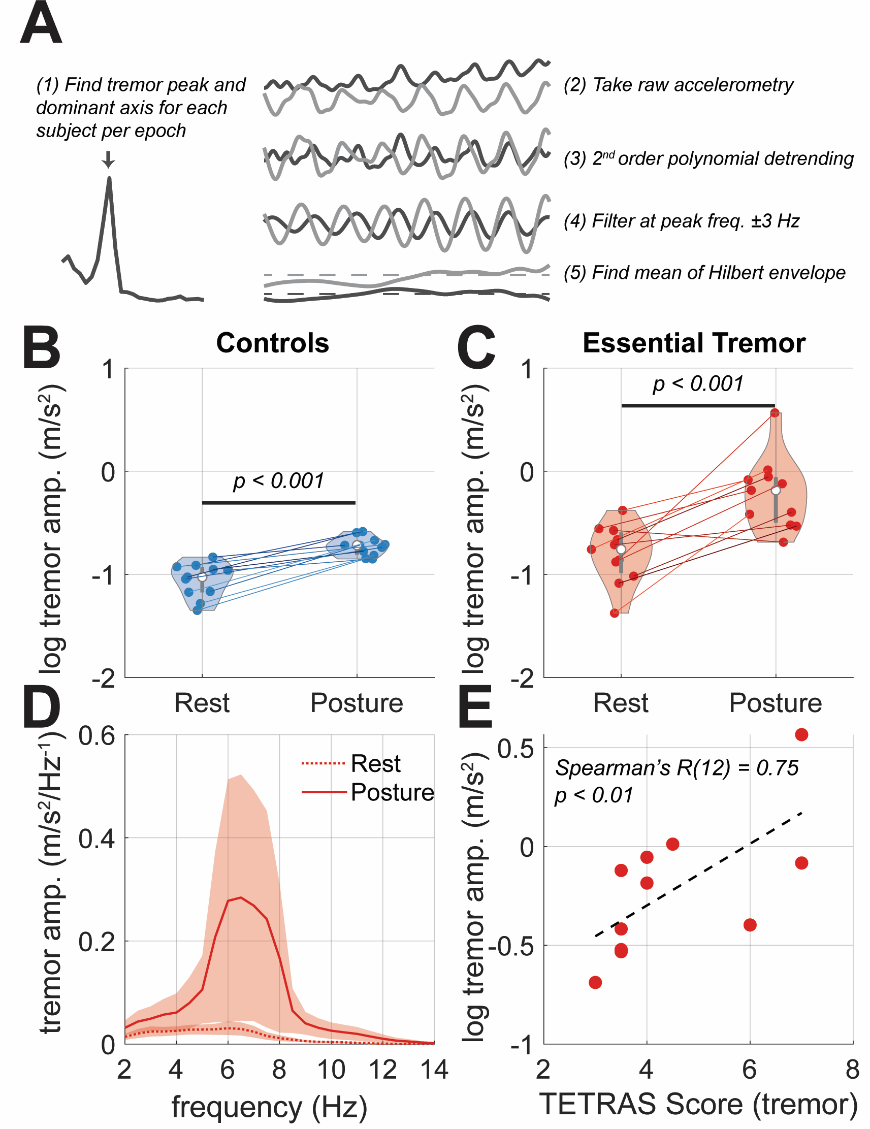

Supplementary Figure 1 – **Analysis of tremor dynamics during steady state recordings at rest and posture.** Violin plots indicate the distribution of tremor amplitudes over either the control (blue) or ET (red) cohorts), with lines linking individuals between rest and posture. Statistics indicate outcomes of paired t-tests. **(A)** Illustration of processing pipeline to compute the tremor amplitude. **(B)** Control subjects show a small increase in physiological tremor from rest to posture. **(C)** As expected, ET subjects show a significant increase in tremor at posture. **(D)** The average tremor peak frequency was around 6 Hz, increasing slightly during posture trials. Bounds give the SEM across ET patients. **(E)** Measures of tremor amplitude at posture positively correlate with clinical TETRAS assessment of tremor severity. A regression line is shown alongside, Spearman’s correlation coefficient.

#### 1.2 Supplementary Figure 2 –Effects of Experimental Conditions on Tremor

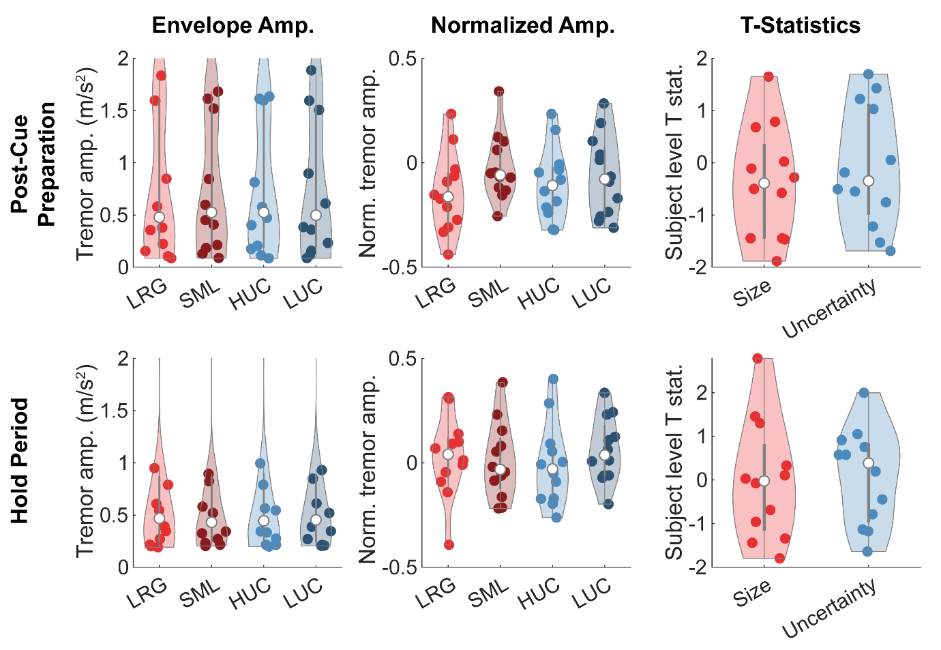

Supplementary Information 2 – **Analysis of modulation of tremor amplitude by task conditions.** Tremor amplitude was estimated as in Figure 2A. We divided trials by the 2x2 conditions modulating either target size (SML or LRG) and the uncertainty of directional cues (HUC or LUC). (1^st^ Column) The violin plots here show the distribution of tremor power across the conditions. Each point represents the mean of an individual subject. (2^nd^ Column) The distributions of tremor amplitude normalized by the subject level mean and standard deviation. (3^rd^ Column) A plot of the subject level T-statistics comparing each of the experimental conditions. No significant differences between conditions were found.

#### 1.3 Supplementary Figure 3 - Source level distribution of movement responsive beta power

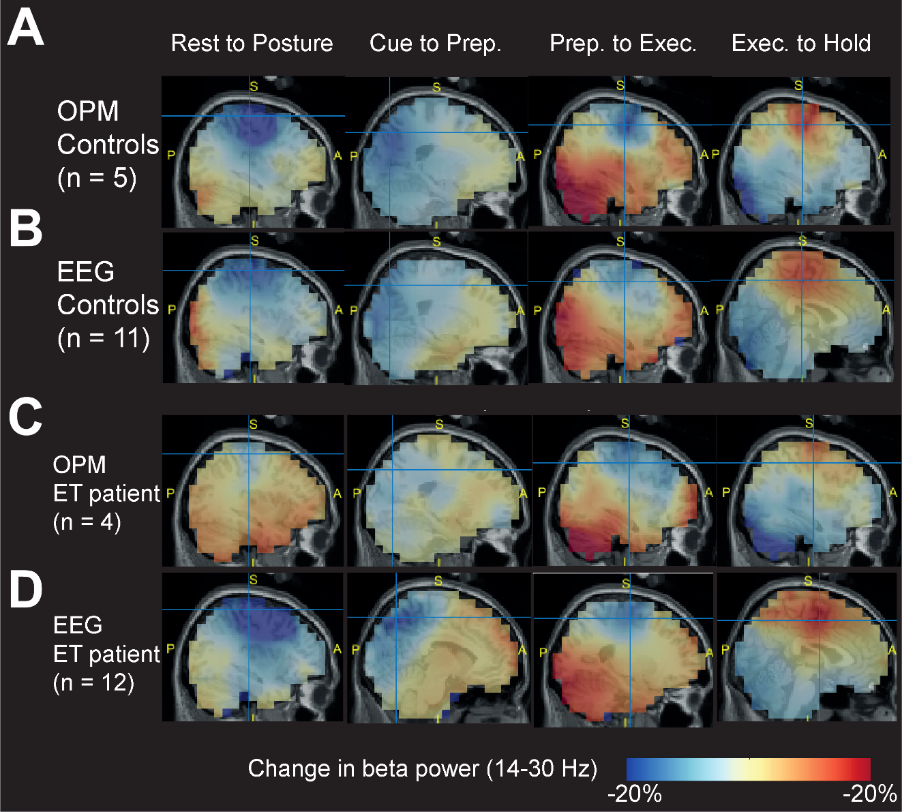

Supplementary Figure 3 – **Source level distribution of 14-30 Hz beta power.** Beta power was localized using a DICS beamformer. Differences in power relative to transitions during the reaching task were computed. Source images were normalized at a subject level before averaging. **(A)** Data for age-matched controls recorded with OPMs; **(B)** age-matched controls recorded with high density EEG; **(C)** Essential Tremor patients recorded with OPMs; **(C)** Essential Tremor patients recorded with high density EEG.

#### 1.4 Supplementary Figure 4 - Spectrograms for Virtual Electrodes in Control Subjects

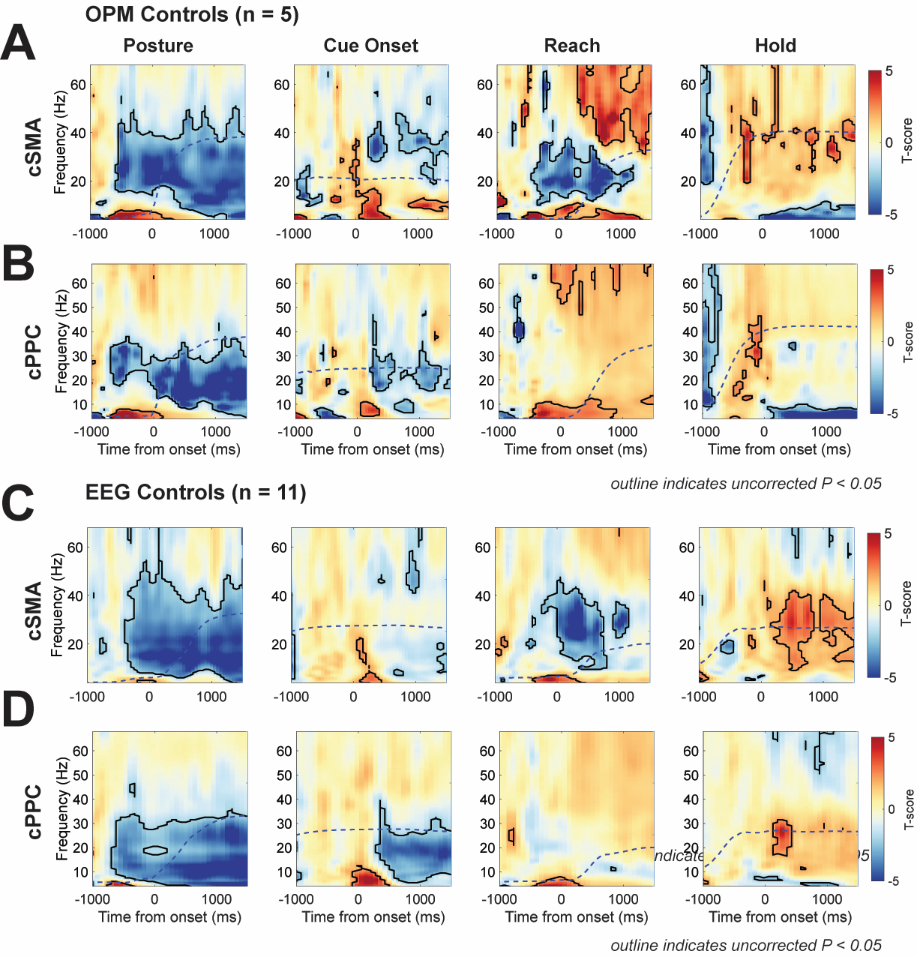

Supplementary Figure 4 - **Time-frequency spectrograms of time locked activity in either the cSMA or cPPC, in both OPM and EEG data in control subjects.** Panels show the group level t-statistics of time-frequency spectrograms constructed from virtual channels, compared to baseline. Bold outlines indicate thresholding on the critical T (P < 0.05; uncorrected). Dashed lines overlaid indicate the group averaged movement trace. **(A)** Spectrograms of OPM derived cSMA activity for time locked activity at onset of postural hold (1^st^ column); onset of directional cues (2^nd^ column); onset of reach (3^rd^ column); and establishment of the hold period (4^th^ column). **(B)** Same as (A), but for cPPC activity. **(C-D)** Same as A-B, but for EEG data.

#### 1.5 Supplementary Figure 5 - Spectrograms for Virtual Electrodes in Essential Tremor Subjects

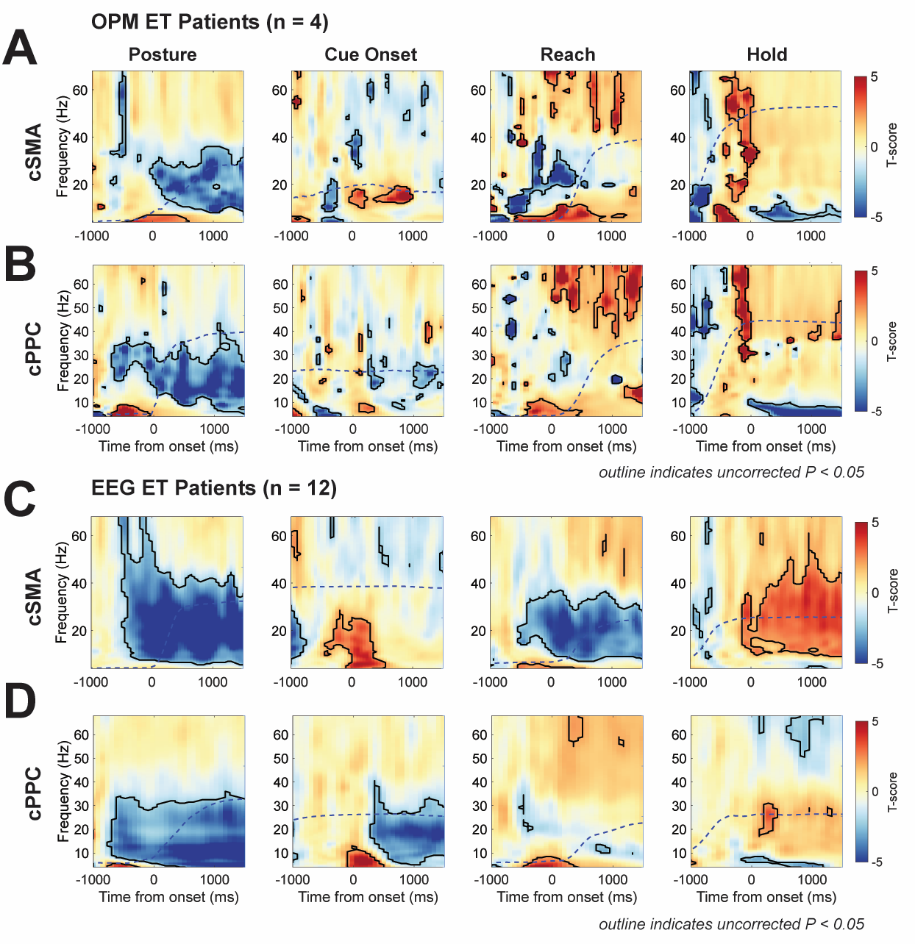

Supplementary Figure 5 – **Time-frequency spectrograms of time locked activity in either the cSMA or cPPC, in both OPM and EEG data in ET patients.** Panels show the group level t-statistics of time-frequency spectrograms constructed from virtual channels, compared to baseline. Bold outlines indicate thresholding on the critical T (P < 0.05; uncorrected). Dashed lines overlaid indicate the group averaged movement trace. **(A)** Spectrograms of OPM derived cSMA activity for time locked activity at onset of postural hold (1^st^ column); onset of directional cues (2^nd^ column); onset of reach (3^rd^ column); and establishment of the hold period (4^th^ column). **(B)** Same as (A), but for cPPC activity. **(C-D)** Same as A-B, but for EEG data.

#### 1.6  Supplementary Figure 6 – Comparison Of Whole Brain, Motor Responsive Latent Networks Between Controls and ET Patients Using OPM Data

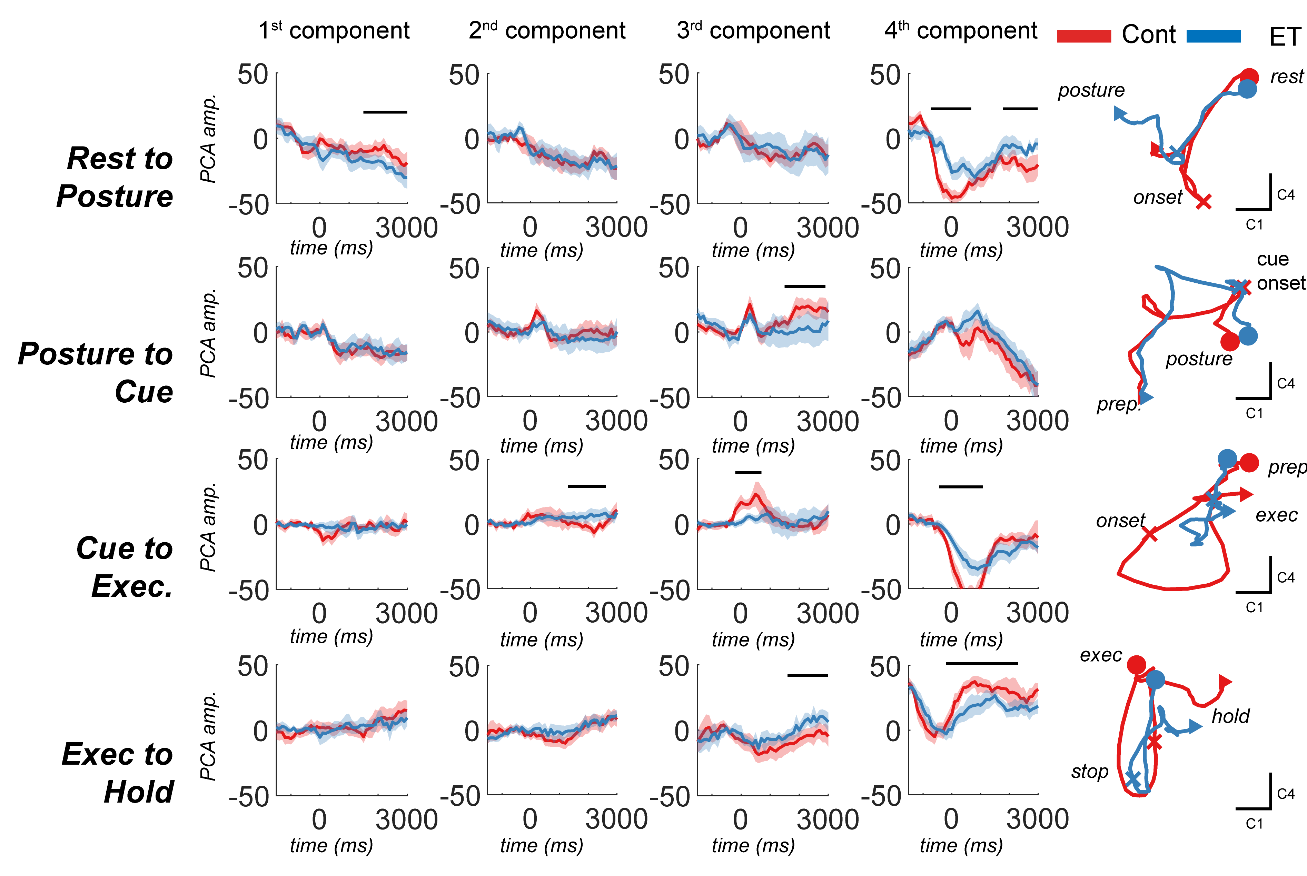

Supplementary Figure 6 –**Visualization of whole brain, motor responsive, latent dynamics and a comparison between controls and patients with ET recorded with OPM.** Components were computed using tfPCA applied to the group averaged EEG data (Figure 5). These coefficients were then used to project data to trial-level latent dynamics that could be used to explore differences between controls and ET patients. **(A)** The latent dynamics exhibited during the postural hold for each component (columns) are indicated for ET (blue) and controls (red) separately. Bars and associated P-values show the outcome of cluster permutation statistics between the two experimental groups. **(B-D)** Same as (A) but for cue presentation, reach execution, and the sustained hold. **(E-H)** Plots of the 2D latent trajectories (components 1 and 4, only) indicate highly correlated dynamics between controls and ETs with quantitative differences in the weighting of the components such as increased engagement of the prefrontal beta network in ET subjects (4^th^ component, apparent in plot G).

#### 1.7  Supplementary Figure 7 – Comparison Of Whole Brain, Motor Responsive Latent Networks Between Controls and ET Patients Using OPM Data

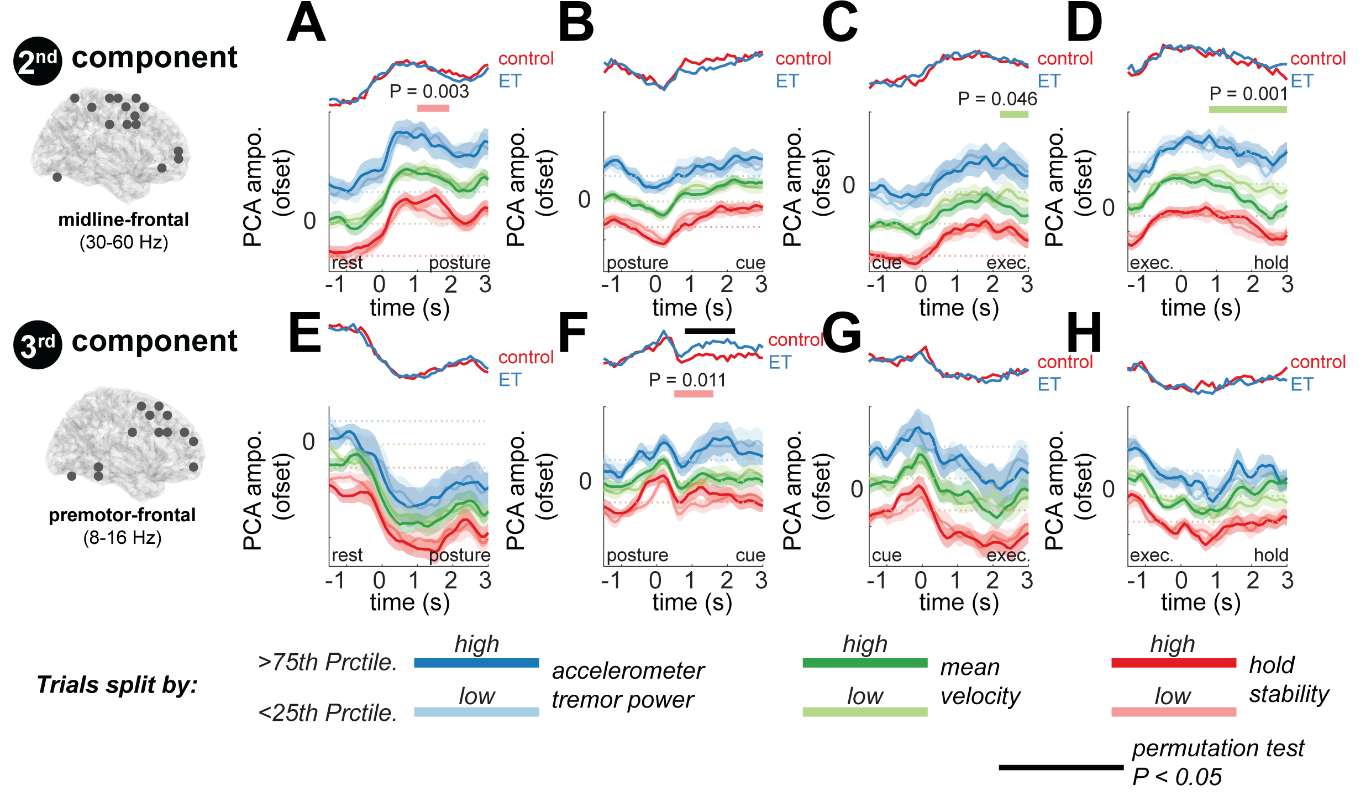

Supplementary Figure 7 –**Visualization of whole brain, motor responsive, latent dynamics and a comparison between controls and patients with ET recorded with OPM.** Components were computed using tfPCA applied to the group averaged EEG data (Figure 5). These coefficients were then used to project data to trial-level latent dynamics that could be used to explore differences between controls and ET patients. **(A)** The latent dynamics exhibited during the postural hold for each component (columns) are indicated for ET (blue) and controls (red) separately. Bars and associated P-values show the outcome of cluster permutation statistics between the two experimental groups. **(B-D)** Same as (A) but for cue presentation, reach execution, and the sustained hold. **(E-H)** Plots of the 2D latent trajectories (components 1 and 4, only) indicate highly correlated dynamics between controls and ETs with quantitative differences in the weighting of the components such as increased engagement of the prefrontal beta network in ET subjects (4^th^ component, apparent in plot G).

#### 1.8 Supplementary Figure 8 – Convolutional Neural Network for Movement Detection

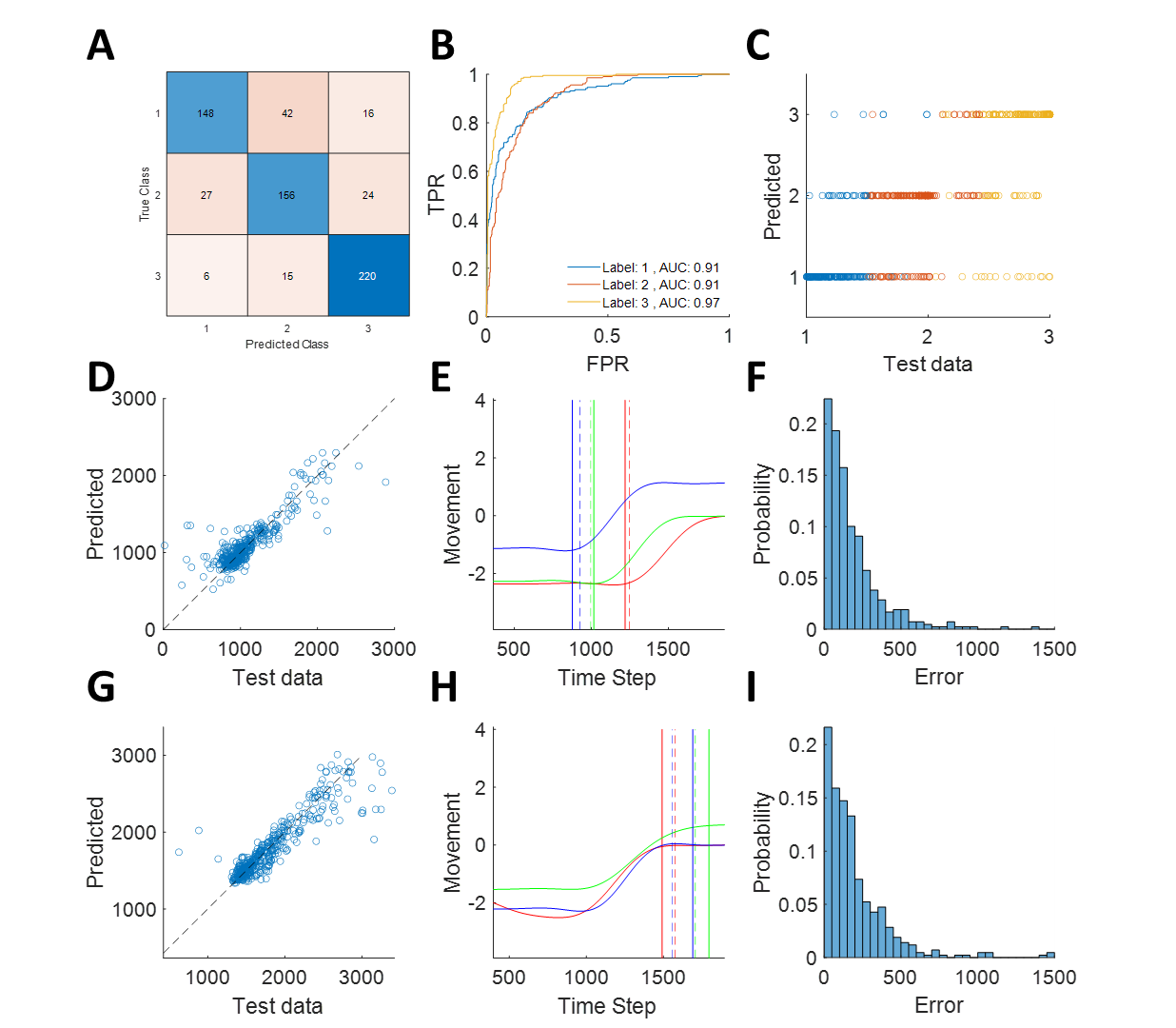

Supplementary Figure 8 – **Validation of convolutional neural network model for time locking of reaching movement.** Plots indicate results examining the performance of a CNN based classifier/regression for reach quality (A-C); reach onset (exec.; D-F) and termination (hold; G-I). Training data was provided by manual labelling of data from three separate assessors on ~30% of the data. **(A)** Confusion matrix of recording quality class: (1) is optimal, (2) sub-optimal, (3) is rejected. See Supplementary Information II for criteria. **(B)** Receiver operating characteristics for reach quality classifier, with each of the three data quality labels. **(C)** Scattergram of predicted vs test data. **(D)** Same as (C) but for prediction of reach onsets. **(E)** Example plot of three predicted (bold) vs manually marked (dashed) reach onsets. **(F)** Histogram of the model error (|predicted-manual|, ms). **(G-I)** Same as D-F, but for reach holds.

### 2 Supplementary Information

#### 2.1  Supplementary Information I – Clinical Table: EEG Patients

| Subject | Age | Gender | TETRAS Performance | TETRAS Upper Limb Tremor (R/L) | TETRAS Head Tremor | Medication |
| --- | --- | --- | --- | --- | --- | --- |
| 1 | 53 | M | 12.5 | 3.5/3 | 0 | Gabapentin 600 mg; Primidone 500mg; Amitriptyline 75 mg |
| 2 | 73 | M | 15.5 | 3/4 | 1 | Primidone 150mg; Propranolol 40mg; L-Thyrpx 100mg; Ramipril 2.5 mg |
| 3 | 68 | M | 23 | 4/6 | 0 | Propranolol 40mg; Amlodipine 5mg; |
| 4 | 59 | M | 38 | 6.5/8 | 1 | Propranolol 80mg; Amlodipine 5mg; Insulin Glargine + Lispro; Clopidogrel 75mg; Simva 20mg; Ramipril 10mg |
| 5 | 19 | F | 7.5 | 1/3.5 | 0 | None |
| 6 | 19 | M | 11 | 3/3 | 0 | None |
| 7 | 58 | F | 15.5 | 3/4 | 1 | Propranolol 10mg; Escitalopram 15mg |
| 8 | 24 | F | 9 | 3.5/3 | 0 | None |
| 9 | 21 | F | 17 | 3.5/7 | 0 | Propranolol 10mg |
| 10 | 61 | M | 27.5 | 7/6 | 2 | Propranolol 40mg; Liskantin 250mg |
| 11 | 42 | M | 14 | 4.5/4.5 | 0 | None |
| 12 | 64 | F | 13 | 3/3.5 | 1 | Propranolol 20mg |

#### 2.2  Supplementary Information II – Clinical Table: OPM Patients

| Subject | Age | Gender | TETRAS Performance | TETRAS Upper Limb Tremor (R/L) | TETRAS Head Tremor | Medication |
| --- | --- | --- | --- | --- | --- | --- |
| 1 | 67 | F | 19 | 5/4 | 0 | None |
| 2 | 44 | F | 25 | 7/7 | 0 | None |
| 3 | 66 | M | 17 | 6/6 | 0 | Amlopidine, Pravastatin, NiC for tremor |
| 4 | 25 | F | 17 | 3/6 |  | Primidone 25 mg |

#### 2.3 Supplementary Information III – Analysis of Reproducibility of Latent Dynamics

**Table 1 –** Analysis of Correlation Between Latent Dynamics in Controls and ET Recordings

|  | **EEG** | | | | |
| --- | --- | --- | --- | --- | --- |
|  | **Comp 1** | **Comp 2** | **Comp 3** | **Comp 4** | *Average* |
| **Posture** | 0.96 | 0.98 | 0.98 | 0.94 | *0.96* |
| **Cue** | 0.91 | 0.95 | 0.68 | 0.86 | *0.85* |
| **Reach** | 0.90 | 0.98 | 0.91 | 0.95 | *0.94* |
| **Hold** | 0.88 | 0.90 | 0.88 | 0.98 | *0.91* |
| **Average** | *0.91* | *0.95* | *0.86* | *0.93* | ***0.91*** |
|  | **OPM** | | | | |
|  | **Comp 1** | **Comp 2** | **Comp 3** | **Comp 4** | *Average* |
| **Posture** | 0.90 | 0.90 | 0.83 | 0.88 | *0.88* |
| **Cue** | 0.96 | 0.68 | 0.42 | 0.97 | *0.76* |
| **Reach** | 0.84 | 0.85 | 0.89 | 0.94 | *0.88* |
| **Hold** | 0.87 | 0.90 | 0.76 | 0.85 | *0.85* |
| *Average* | *0.89* | *0.83* | *0.73* | *0.91* | ***0.84*** |

**Table 2 –** Analysis of Correlation Between Latent Dynamics in OPM and EEG

| Analysis of Correlation Between Latent Dynamics in OPM and EEG (Pearson’s Correlation Coefficient R) | | | | | |
| --- | --- | --- | --- | --- | --- |
|  | **Component 1** | **Component 2** | **Component 3** | **Component 4** | *Average* |
| **Posture** | 0.93 | 0.72 | 0.65 | 0.93 | *0.81* |
| **Cue** | 0.94 | 0.62 | 0.47 | 0.99 | *0.76* |
| **Reach** | 0.97 | -0.03 | 0.46 | 0.94 | *0.59* |
| **Hold** | 0.93 | 0.56 | 0.65 | 0.87 | *0.75* |
| *Average* | *0.94* | *0.47* | *0.56* | *0.93* | ***0.73*** |

#### 2.4  Supplementary Information IV – Algorithm to Manipulate Cue Uncertainty

Let *D* be the set of cardinal directions, where *D = {N, NE, E, SE, S, SW, W, NW}*. A direction *d* is selected from *D*, where *d* ∈ *D*. We generate a set of N random phases (expressed in radians):

$$\theta_{n}=\bar{\theta}+\sigma\epsilon_{n}, n=1,2,\ldots,N$$

where $\bar{\theta}$and $\sigma$ represent the chosen angle and dispersion (uncertainty) of the movement cues, and $\epsilon_{n}$ is a random variable sampled from a standard normal distribution, $\epsilon_{n}\sim N(0,1)$. The direction of each arrow is then wrapped around the unit circle: $\phi_{n}=\arg\left( e^{i\theta_{n}} \right).$The arrow directions are then determined by computing the normalized histogram for bins with edges corresponding to the cardinal directions, to yield a scalar for each arrow. We used $\sigma$= 0.98 for low uncertainty trials, and $\sigma$= 1.64 for high uncertainty trials, yielding 85% and 15% average success rates in piloting with healthy controls.

#### 2.5  Supplementary Information V – Criteria for Reach Quality Marking

*Grade 1 – Optimal Reach*

- A stable baseline is present prior to movement, with little oscillation or prepotent movement.
- The reach may be initiated before the movement cue (i.e., t < 0) if baseline is stable.
- The reach itself is smooth, with a clear stereotyped trajectory.
- The hold period is stable and lasts greater than 2 seconds, with no clear drifts or repositioning of the hand.

*Grade 2 – Suboptimal Reach*

- A baseline period is apparent, although some slow drifts in posture, or small prepotent movements are present.
- The reach may be initiated before the movement cue (i.e., t < 0) if the baseline is stable.
- The reach is jerky, or non-smooth containing mid-reach corrections and changes in velocity.
- The hold period is apparent and last for a duration >= 2 seconds. The hold period may contain slow drifts, or small corrective movements.

*Grade 3 – Rejected Reach*

- The reach is slow (> 4 seconds).
- The reach is jerky/non-smooth.
- There is no clear baseline periods lasting > 2 seconds.
- There is no clear hold period lasting > 2 seconds.

#### 2.6  Supplementary Information VI – Details of Source Estimation Techniques

Source inversion of sensor-level data involves constructing forward models to simulate the propagation of neural activity from the source to the sensor. For EEG data, we used a template boundary element model with sensor locations set according to the 10-10 system. For OPM data, sensor positions were derived from custom head cast models aligned to the subject’s structural MRIs. For the two subjects lacking MRIs, sensors were transformed to a template space using an iterative closest point algorithm to match the 3D scanned scalp to the template model. OPM head models used the single shell method^103^. To facilitate group-level statistical comparisons, a nonlinear warping was applied to align the anatomical scans.

Subject level common filters were computed from data concatenated across the multiple epochs of the experiment. Covariance matrices were computed in two frequency bands: tremor (peak postural tremor frequency ±1.5 Hz) and wide band beta (14-30 Hz). Each covariance matrix was truncated by its effective rank^104^. A covariance regularization of 1% was used.

#### 2.7  Supplementary Information VII – Implementation of Time Frequency Principal Component Analysis

tfPCA was applied to the group-averaged data with spectrograms concatenated for each motor epoch (time x four epochs). To avoid low frequency bias due to 1/f structure of EEG/OPM, spectra were log scaled. Spectrograms were also Z-normalized per subject to remove differences in SNR. We then constructed an *n × p* matrix ***X*** ([*n*: concatenated time] x [*p*: channel x frequency]) to which we applied PCA to yield coefficients ***V*** (*p* x number of components) for each spatio-spectral component, and latent scores ***Z*** (component x time). PCA coefficients were rotated using the Varimax algorithm to aid interpretability. We then projected components from the single trial level data *D_t_* (to compare experimental conditions and perform regressions) by computing *Z_t_ =* ***V****D_t_*. We also reconstructed components in the time frequency domain by multiplying trial-level data with the projection matrix ***P = VV^T^***, which we used for CNN tremor prediction analyses. In this way, the coefficients derived from the group averaged EEG data were used to project both EEG and OPM data to single trial level components.

#### 2.8 Supplementary Information VIII – Details of 3^rd^ Party Toolboxes

| Toolbox Name | Author | Year | Source/Reference |
| --- | --- | --- | --- |
| boundedline-pkg | Kelly Kearney | 2015 | <https://github.com/kakearney/boundedline-pkg> |
| brewermap | Stephen Cobeldick | 2014 | <https://github.com/DrosteEffect/BrewerMap> |
| Fieldtrip | Donders Institute, Radbound University | 2020 | <https://www.fieldtriptoolbox.org/> |
| linspecer | Jonathan C. Lansey | 2015 | <https://github.com/davidkun/linspecer> |
| Matlab Toolbox for Dimensionality Reduction (v0.8.1b) | Laurens van der Maaten | 2013 | <https://lvdmaaten.github.io/drtoolbox/> |
| splitvec | Bruno Luong | 2009 | [ttps://uk.mathworks.com/matlabcentral/fileexchange/24255-splitvec](https://uk.mathworks.com/matlabcentral/fileexchange/24255-splitvec) |
| SPM 12 | Wellcome Centre for Human Neuroimaging, University College London | 2020 | <https://www.fil.ion.ucl.ac.uk/spm/> |
